# When a look means nothing: contextual interpretation abolishes the Gaze Cueing Effect

**DOI:** 10.1101/2024.07.18.603902

**Authors:** Amit Zehngut, Shlomit Yuval-Greenberg

## Abstract

In a world overloaded with sensory input, organisms use spatial attention to selectively process the most relevant information. For a highly social species like humans, one influential attentional cue is the direction of other people’s gaze: by perceiving where others look and following their gaze, observers can increase their chances of detecting important information. This well-established phenomenon, called the *gaze cueing effect* (GCE), is traditionally considered reflexive. However, gaze-shifts do not always indicate external attention; they can also occur for other reasons, such as when attention is shifted inwards during effortful cognitive processing. The ability of observers to interpret the eye movements of other people correctly, while dissociating inwards and outwards attention shifts, is critical for efficient management of their attentional resources.

In this study, we examined the role of context on the interpretation of other people’s gaze. Across two preregistered experiments (total *N*=110), participants viewed gaze-shifts while performing a perceptual task. One group was primed to interpret gaze-shifts as reflecting inward attention deployment during cognitive effort, while the other was not provided with an interpretive context. The results revealed that GCE was reduced when gaze-shifts were perceived as linked to cognitive effort rather than external attention, indicating that attentional responses to gaze cues are context-dependent rather than purely reflexive. These findings highlight the flexibility of social attention, revealing that higher-order cognitive interpretation can override well-established attentional mechanisms. With its interdisciplinary approach, reexamining attentional processes in light of social cognition, this study offers new perspectives on the multifaceted socio-cognitive mind.

**Significance statement:** In a world overflowing with sensory input, our attention is a limited resource which must be allocated efficiently. One way we manage this is by attending to locations where other people look, assuming their gaze points to something important. This study shows this mechanism of attention shift is not automatic. Instead, individuals interpret what the eye movements reflect and modulate their attention shifts accordingly. When observers believed that a person was thinking rather than looking at something meaningful, they were less likely to shift their attention in the same direction. These findings challenge the long-standing idea that humans reflexively follow others’ gaze and reveal that our attention is guided not just by where people look, but by what we think is happening in their minds. This insight has real-world implications for improving communication, education, and the design of socially aware technologies like virtual assistants or classroom tools.

## Introduction

In a world overloaded with sensory input, humans and other animals rely on spatial attention to selectively process information. By prioritizing certain regions of the environment, organisms can efficiently allocate cognitive resources to what matters most. This selective process may occur *exogenously*, triggered by external salient stimuli, or *endogenously*, guided by cues indicating where a target is most likely to appear. For a highly social species such as humans, one of the most influential endogenous cues is the direction of other people’s gaze: by perceiving where others look and follow their gaze direction, observers can increase their chances of detecting important information in the environment, while conserving their own attentional resources.

A classic demonstration of how people align their attention with another person’s gaze is the *gaze cueing effect (GCE)*. When individuals witness someone else shift their gaze, their own attention often shifts in the same direction, speeding their response to targets located at the observed gaze trajectory, relative to targets located elsewhere (Driver et al., 1999; Friesen & Kingstone, 1998). This phenomenon highlights a social dimension of spatial attention: we routinely follow other people’s gaze directions, presumably because – across evolutionary development – following someone else’s gaze often led to survival benefits (Frischen, Bayliss & Tipper, 2007). The GCE is typically rapid, unconscious and involuntary, and was therefore often considered to be reflexive (Driver et al., 1999; Friesen, Moore, & Kingstone, 2005; McKay et al., 2021; Ristic, Wright & Kingstone, 2007; Schmitz et al., 2024). However, similarly to other reflexive behaviors, the GCE can be modulated by top-down factors, such as the observer’s characteristics (e.g. gender, age, social status and mental state) and the situational context (Bayliss, Di Pellegrino & Tipper, 2005; Bayliss, Schuch & Tipper, 2010; Frischen & Tipper, 2006; Slessor et al., 2016; Teufel, et al., 2010; Dalmaso, Castelli & Galfano, 2020). Moreover, GCE can be induced by contextual interpretation. When symbolic items are primed to be perceived as social cues, they can trigger a GCE (Schmitz & Einhäuser, 2023).

The evolutionary perspective on the GCE suggests that following someone else’s gaze enhances survival prospects by directing attention to potentially crucial information, assuming that another person’s gaze shift signifies their priority selection. But is this an accurate assumption? Do gaze shifts invariably correspond to attentional shifts and prioritization? The answer is no. Despite the well-known link between eye movements and attention shifts, there are instances where gaze may be directed to non-informative or irrelevant locations. For example, the ‘looking-at-nothing’ effect shows that people retrieving memorized information often shift their gaze to empty locations where the information was previously located, even if it is no longer there (Johansson & Johansson, 2014; Taub & Yuval-Greenberg, 2023). Another example is *gaze aversion (GA)*; during cognitively demanding tasks, like formulating a response to a question, individuals tend to avert their gaze away from a conversational partner onto the far periphery (Doherty-Sneddon, 2002). These large eye movements are thought to reduce external distractions while focusing attention inward during demanding cognitive processing (Doherty-Sneddon & Phelps, 2005; Abeles & Yuval-Greenberg, 2017, 2021). Notably, GAs do not reflect spatial attention orientation, with their target locations being uninformative, so orienting one’s own attention in response to other people’s gaze aversions carries no attentional benefit.

These observations raise an important question: How do observers determine whether another person’s gaze signals a location worth attending to, or merely reflects internal cognitive processing? What factors lead certain eye movements to serve as spatial gaze cues, while others are perceived as attentionally uninformative gaze aversions? One possible answer is that observers rely on kinematic properties of an eye movement (e.g. its velocity, direction or eccentricity) to infer whether this eye movement indicates attentional shift or internal processing. However, evidence relating certain kinematic characteristics of eye movements to their attributed attentional role is ambivalent (Servais, Hurter & Barbeau, 2022, Friesen, Ristic & Kingstone, 2004). Here we hypothesize that context – rather than movement metrics – shapes the interpretation of observed gaze shifts. We further propose that the *same* eye movement, with the *same* kinematic attributes, can either be perceived as directing attention externally or signifying internal thought, depending on the cognitive state attributed to the gazing individual.

To test this idea, we conducted two pre-registered experiments in which participants viewed identical gaze shifts, followed by a peripheral letter discrimination task. Crucially, participants of one group were led to believe that the gazing individual was performing arithmetic calculations (*Thinking* context), and participants of another group were given no such context (*Attention* context). We predicted that participants lacking additional context would show the classic GCE, while participants informed about the target’s internally focused thought process would not exhibit this effect. Finding this would underscore a dynamic model of social gaze cueing, where contextual framing – not just reflexive responses to eye movements – governs whether observers treat a gaze shift as an attentional cue.

## Methods

### Participants

A total of 122 healthy individuals participated in this study. Participants were randomly assigned to one of the two groups (“Thinking” or “Attention”). We applied two pre-registered exclusion criteria. First, participants were excluded from the study if they reported understanding that the goal of the thinking manipulation was to modulate the way the cueing face is interpreted or if they reported understanding any other crucial aspect of the manipulation. A total of three participants were excluded due to this reason from **Experiment 1** and one was excluded from **Experiment 2**. In addition, since this was an easy task, participants who performed the task with less than 80% accuracy were assumed to have not been engaged in the task and were excluded from the analysis. This has led to the exclusion of seven participants in **Experiment 1** and one in **Experiment 2**. The remaining participants included **50** in **Experiment 1** – 25 in the Attention group (14 females, 2 left-handed, mean age 23.7 ± 1.9 SD) and 25 in the Thinking group (17 females, 5 left-handed, mean age 24.0 ± 3.0 SD); and **60** in **Experiment 2** – 30 in the Attention group (23 females, 2 left-handed, mean age 23.4 ± 2.6 SD) and 30 in the Thinking group (27 females, all right-handed, mean age 23.1± 2.3 SD). Sample size was determined according to a pilot study consisting of 42 participants (21 in each group). A power simulation based on this pilot study indicated that with a sample size of 50 participants, the interaction effect could be observed with a power (1-β) of approximately 0.8. All participants reported having normal or corrected-to-normal vision and no history of neurological or psychiatric disorders. Participants received payment or course credit for their participation. All participants signed a consent form before participating. The experiment has been approved by the ethical committees of Tel-Aviv University and the School of Psychological Sciences at Tel-Aviv University.

### Stimuli

All stimuli were displayed on a gray background (RGB: [0.75,0.75,0.75]). The fixation point was a white cross, measuring 1.3° wide and 1.3° high, positioned 3.4° below the center of the screen (degree symbols ° represent visual degrees). The cue display consisted of a brief video of an individual shifting his gaze, in parallel with an auditory stimulus. The filmed individual (male, age 31) was positioned on the screen such that the fixation point was located between his eyes (approximately 0.3° from each eye). The auditory stimulus was numbers or an arithmetic question read-out in Hebrew by a male speaker. The target stimulus was a black capital letter L or T, measuring 0.8° wide and 1.3° high. In **Experiment 1**, the target was located 10.7° from fixation (3.4° on the vertical axis, and 10.1° on the horizontal axis to the right or left according to target position). Since some of the participants in **Experiment 1** reported perceiving the cuing gaze as directed above the target rather than at the target, we have shifted the target’s location in **Experiment 2** to be slightly upward from the horizontal midline. Therefore, the target was located at an eccentricity of 12.6° from the center of the screen, 7.6° on the vertical axis, and 10.1° on the horizontal axis.

### Experimental Design

Participants were seated in a dimly lit room, with a computer monitor placed 100 cm in front of them (24-inch LCD ASUS VG248QE, 1920×1080 pixels resolution, 60 Hz refresh rate). During the experiment, participants rested their heads on a chinrest. The monitor was adjusted such that the participants’ eyes were positioned at the same height as the fixation point. The experimental design and stimuli were generated using PsychoPy software (Peirce et al., 2019).

Each trial began with a fixation display presented for 500ms. Then a video (“cueing video”) depicting an individual was played for approximately 3.5 seconds, in parallel with an auditory recording. The depicted individual was gazing straight ahead at the camera until slightly before the end of the video (approximately 100ms), when he either continued looking straight (neutral trial, 33% of the trials) or shifted his gaze up-left or up-right (valid and invalid trials, each 33% of the trials). Participants of the “Thinking” group listened to a recording of an arithmetic question that was played simultaneously with the video. This recording consisted of a two-digit number, an operator (–, +, or ×), and a second two-digit number (e.g., “54+91”). The purpose was that the presentation of an arithmetic question in parallel to the video would lead the participants to interpret the situation as an arithmetic question being read-out to the filmed person, which will then think about the answer. Participants of the “Attention” group listened to a recording of a sequence of three numbers: a two-digit number, a single-digit number, and another two-digit number (e.g., “54,1,91”), also played simultaneously to the video. Thus, this group was not led to believe that the situation is of a person thinking of an answer.

At the beginning of the experiment, participants received instructions. In **Experiment 1**, participants of the “Thinking” group were told that in each trial they would see a person listening to arithmetic problems and attempting to solve them and that the targets would appear as soon as the person begins to think of the solution. Participants of the “Attention” group were told that they would see a person and hear a sequence of numbers. In **Experiment 2**, none of the groups received these instructions. In addition, in both experiments and for both groups participants were told that the video was task-irrelevant and were asked to ignore it.

In **Experiment 1**, the video froze at its end for an additional fixed 500ms (an interval which served as a *stimulus onset asynchrony*; SOA) depicting the individual gazing upward left or right, or looking straight, in different trials. In **Experiment 2**, this SOA was modified to be jittered such that the same frame froze for 500±100ms (400-600ms, 10 randomized steps with a step size of 20ms) instead of a fixed interval. This modification was intended to prevent anticipatory responses.

Following the cuing video, the target letter (T or L) was presented at 12.6° up-right or up-left of fixation. Third of the trials were ‘valid’ trials, in which the target was presented on the same side as the direction of the gaze shift. Another third of the trials were ‘invalid’ trials in which the target was presented on the opposite side. The remaining third of the trials were ‘neutral’ trials, where the gazing individual kept looking straight, without any gaze-cue. The target remained on screen until response, but no longer than two seconds. Trials in which no response was detected within two seconds were considered incorrect. Participants were requested to indicate the identity of the target letter by pressing one of two keys labeled “L” or “T”. These labels were applied to the keys “z” (positioned left) and “/” (positioned right) with colored stickers and were counterbalanced between participants. For participants in the “Attention” group, this was the end of the trial and the next trial began immediately after. Participants of the “Thinking” group were presented with another video lasting 1-4 seconds, in which the depicted individual shifted his gaze back to the center and then announced the correct solution for the arithmetic question. Then the trial ended, and the next trial began. The trial procedure is depicted in Figure 1.

**Figure 1.**
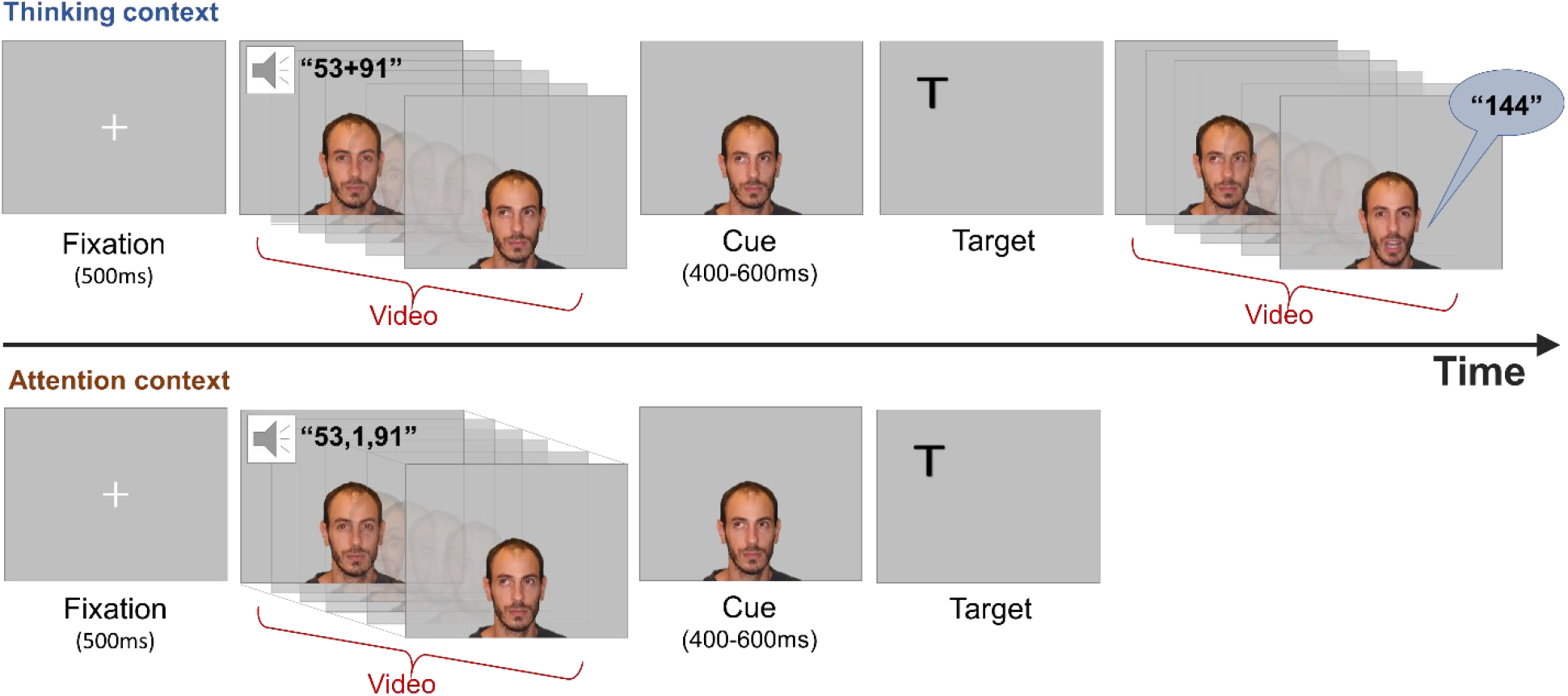
*Trial procedure in each condition*. Each trial began with a fixation display for 500ms. Then a video depicting an individual (the first author) was played for 3.5s, in parallel with the auditory recording. The depicted individual was gazing straight until the end of the video, when he either continued looking straight (neutral trial) or shift his gaze up-left or up-right (valid and invalid trials). The film froze at its end for an additional 500ms (in **Experiment 1**) or 500±100ms (in **Experiment 2**) depicting the gaze cue.Following this, a target letter was presented right or left of fixation. Participants were requested to indicate the identity of the target letter by pressing one of two designated keys. Following target-offset the next trial began, only for participants in the “Attention” group. Participants of the “Thinking” group were then presented with another video, in which the depicted individual shifted his gaze back to the center and then read-out the correct solution for the arithmetic question

For both groups, the experiment consisted of 270 trials in total, with 45 different auditory stimuli, repeating six times each. These included 45 arithmetic questions and their answers in the “Thinking” group and 45 sequences of numbers in the “Attention” group. Cue-validity (valid/invalid/neutral), Gaze direction (left/right/center), Target location (left/right) and Target identity (L/T) were selected at equal probabilities and in pseudo random order.

At the end of each experimental session, participants were debriefed by completing a questionnaire in which they were asked what they assumed to be the purpose of the experiment. Participants of the “Thinking” group were additionally asked to rate the filmed person’s concentration in solving the questions (from 1 to 5), and the percentage of his correct answers (0%, 25%, 50%, 75% or 100%), in their estimation. Participants of the “Attention” group were asked about the task of the filmed person and the reason for playing the sequences of numbers. This questionnaire was used to validate the thinking manipulation.

### Summary of Key Differences Between Experiment 1 and Experiment 2

The preregistered **Experiment 2** was nearly identical to Experiment 1, with a few modifications. First, the target was positioned slightly higher on the screen to align better with the perceived trajectory of the actor’s gaze, addressing participant feedback that the original target location appeared slightly too low. Second, while Experiment 1 provided both groups with some information about the video content, Experiment 2 **omitted all prior information** for both groups. Instead, participants were instructed that the video was **task-irrelevant** and should be ignored, allowing us to examine whether attentional modulation could be based on **implicit, inferable context** rather than explicit instruction. Third, Experiment 1 used a fixed stimulus-onset asynchrony (SOA) of 500 ms, which may have inadvertently induced temporal attention at the cue moment, potentially reducing the GCE. To mitigate this, Experiment 2 introduced a randomized SOA of 400–600 ms.Lastly, to increase statistical power, 60 participants were recruited for Experiment 2, compared to 50 in Experiment 1, with equal distribution between groups.

### Statistical analysis

Analysis followed our pre-registered plan. Trials with incorrect or no response (error trials) or with RT below 150ms, considered unlikely to represent a genuine target-related response (e.g., Keele & Posner, 1968), were discarded from the analysis (3.7% of all trials, range 0–16.3% of trials per participant).

The single-trial RTs of the remaining trials were modeled using a generalized linear mixed model (GLMM), assuming a gamma family of responses with an identity link (Lo & Andrews, 2015).Particularly, this analysis assumes that the RTs follow a gamma distribution, and that the predictors are linearly related to the predicted RT, thus an identity link was used (i.e., no transformation was made on the value produced by the predictors; Lo & Andrews, 2015). The fixed effects were modeled as follows: (1) cue-validity (valid/invalid/neutral) as a within-subject term, to model the GCE; (2) context-group (“Thinking” / “Attention”) as a between-subject term, to model the effect of the manipulation; (3) the interaction terms between group and cue-validity to model the effect of the context on the GCE. Statistical significance for main effects and interaction was determined via a likelihood ratio (LR) test against a reduced nested model, excluding the fixed term. Statistical significance for the parameter coefficients was determined according to the Wald Z test (Fox, 2016).

In addition to the fixed effects, we considered the z-scaled current trial number (i.e., the running trial identifier for the given session) as a covariate to capture the effects of fatigue and training during the experiment (Baayen & Milin, 2010). Since the different experimental groups may have experienced different fatigue or training effects, we additionally considered the interaction between group and trial number. In addition, RTs may be affected by the congruency between the target location (T/L in left or right) and the order of the key positions (T/L in left or right), a phenomenon known as the “Simon effect” (Simon, 1990), therefore we considered the interaction between the target location and the key position.

The random effect structure of the model followed a procedure suggested by Bates et al. (2015). We started with the simplest model, a random intercept-by-subject-only model, and then added a random slope for the within-subject factor cue-validity. We examined whether the new model converges, and then used the likelihood ratio test (using α = .05) between the new and the simple model, to examine whether there is an improved fit over the previous model and avoid overfitting. Next, we tested whether the model supports random slopes for the current trial covariate. It appeared that in each experiment, the full model could not be converged with all covariates and random terms. In both experiments, the full model included all the fixed terms and a random intercept by subject. In **Experiment 1**, the model also included the z-scaled trial number and the interaction between the target location and key position (“Simon effect”) as covariates. In **Experiment 2**, the model included the interaction between trial number and context-group as an additional covariate. The full model selection process is described in the R markdown file in the project OSF repository (see below, Data availability).

For simplicity and consistency with existing research (e.g., McKay et al., 2021), we performed an additional analysis. Mean RTs for correct responses in each of the groups were subjected to a separate one-way repeated-measures ANOVA with cue-validity (valid, invalid, and neutral) as a within-participant factor. Follow-up planned contrasts between valid and invalid levels were applied to measure the effect size of the GCE for each group. The p-values were corrected for multiple comparisons, using a false-discovery-rate (FDR) correction (Benjamini & Hochberg, 1995). To examine whether the results of each ANOVA provide support for null hypothesis (H_0_; i.e., no GCE) compared to the alternative hypothesis (H_1_; i.e., a GCE), we performed a Bayesian analysis for each ANOVA, with an ultrawide Cauchy distribution (scale of effect r = 1). We considered BF_10_>3 and BF_10_< 0.33 to provide conclusive evidence against or in favor H_0_, respectively, and 0.33<BF_10_<3 as inconclusive (Dienes & Mclatchie, 2018; with a BF_10_ between 3 and 10, 10 and 30, 30 and 100 and > 100 providing moderate, strong, very strong, and extreme evidence, respectively, for H_1_; Jeffreys,1961; Wagenmakers et al., 2018). The calculated effect sizes are Cohen’s d standardized repeated-measures mean differences with Hedges’s g corrections (Lakens, 2013). Based on the previous findings, the minimum effect size that has been reported to demonstrate the GCE generally ranges from approximately 0.05 to 0.20 (McKay et al., 2021). Smaller effect sizes (e.g., below 0.05) being less consistently replicated or more likely to reflect marginal effects. To allow comparison between effect sizes, a 95% bootstrapped confidence interval (CI) with 2000 iterations was computed for each effect size. Analyses were performed in R version 4.3.2 using RStudio (https://www.r-project.org). GLMM modeling was performed using the lme4 (Bates et al., 2015) package,Bayesian modeling was performed using the BayesFactor package (Morey & Rouder, 2015).

### Data Availability

The datasets generated by this study and an R markdown file that reproduces all the reported modeling, statistical analyses, and graphs in the article as well as the pre-registered plan, are uploaded to the Open Science Foundation repository and are available at https://osf.io/EGFK7.

## Results

### Experiment 1

We found a significant main effect for cue-validity (χ^2^(4) = 32.4, p < 0.001): RTs were faster to targets located congruently with the actor’s gaze direction, compared to trials in which the actor was gazing straight ahead (valid < neutral; β = 9.8ms, t = 2.14, p = 0.03) and compared to trials in which the actor’s gaze was directed incongruently with the direction of the target (valid < invalid; β = 11.1ms, t = 2.4, p = 0.02). This finding is consistent with the GCE. In addition, we found an unpredicted significant main effect for context (χ^2^(3) = 8.0, p = 0.04), originating from faster RTs in the Attention relative to the Thinking group (β = -91.3ms, t = -2.38, p = 0.017). This main effect was likely due to the tendency of some participants of the Thinking group to calculate the arithmetic problem presented to them, even though they were asked to ignore them. Performing a calculation during the trial, could have slowed the response to the target. No significant interaction was found between group and cue-validity, by estimating the unique contribution of the interaction fixed term to the model (χ^2^(2) = 3.17, p = 0.21). However, two pre-registered simplified ANOVA conducted separately within each group supported the hypothesis by revealing a GCE only in the Attention group (F(2,6460)=8.73, p < 0.001) but not in the Thinking group (F(2,6496)=1.41, p = 0.24). The inconsistency between the ANOVA and the GLMM analyses with regards to the interaction, suggests limited power in the sample of Experiment 1, which was indeed found to contain high interindividual variability in RTs (see Figure 2). Nevertheless, the significant interaction was further supported by a Bayesian analysis suggesting strong evidence for a GCE in the Attention group (BF_10_=10.84) and extreme evidence for no GCE in the Thinking group (BF_10_=0.007). More specifically, In the Attention group, RTs in valid trials were found to be significantly faster compared to invalid (M_Δ_ = 16.6ms, SE = 5.76, t(6460) = 2.87, p = 0.006, Hedges’s g = 0.09; 95% CI for the effect size = [0.03, 0.15]) and compared to neutral trials (M_Δ_ = 23.3ms, SE = 5.74, t(6460) = 4.06, p < 0.001, Hedges’s g = 0.12; 95% CI for the effect size = [0.06, 0.18]). However, in the Thinking group, no significant difference was found between valid and invalid trials (p = 0.28, Hedges’s g = 0.05; 95% CI for the effect size = [-0.01, 0.11]) nor between valid and neutral trials (p = 0.49, Hedges’s g = 0.02; 95% CI for the effect size = [-0.04, 0.08]).

**Figure 2.**
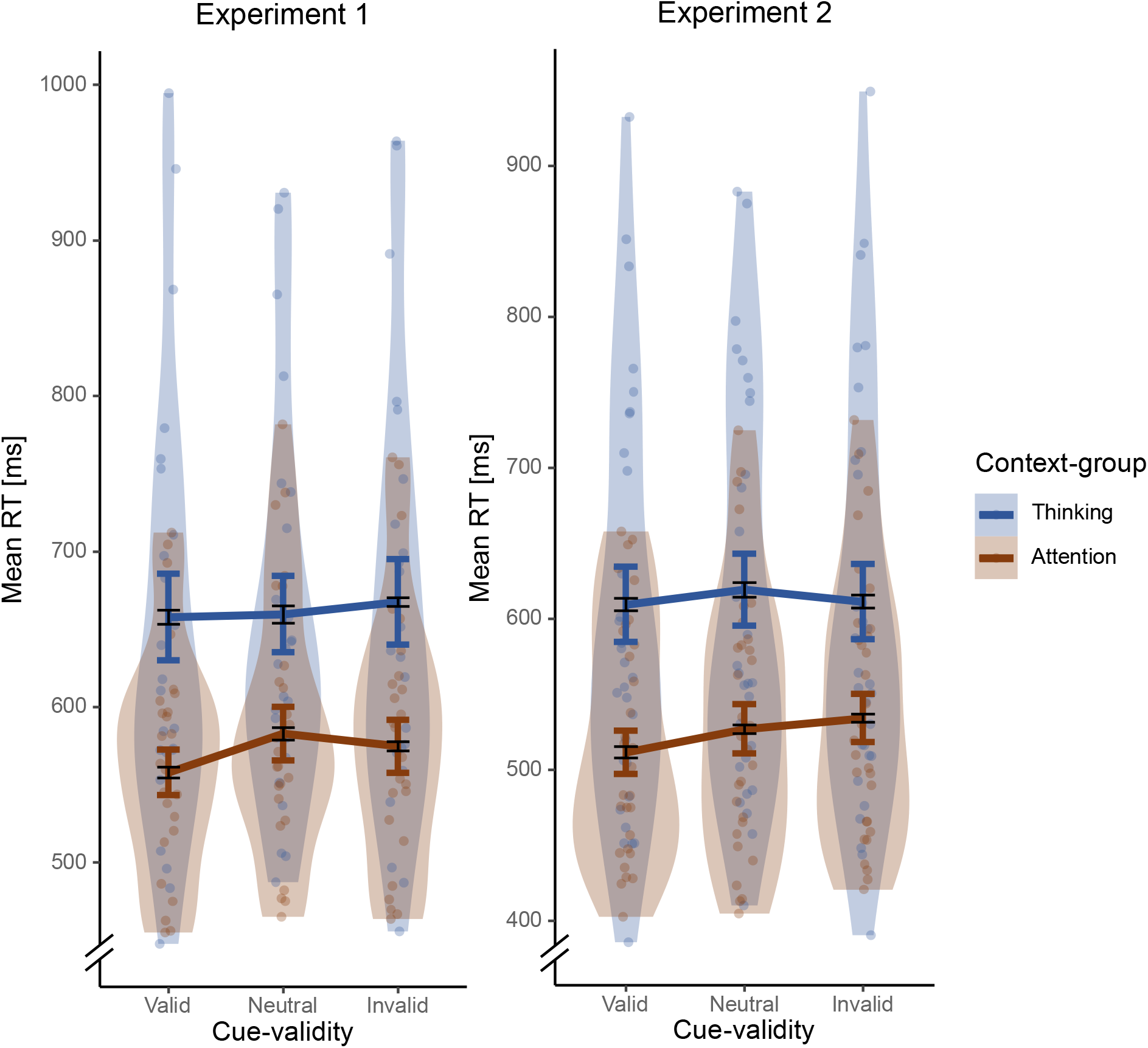
*RTs as a function of cue-validity for each context group*, in **Experiment 1** (left; N=50) and **Experiment 2** (right; N=60). Mean RT for the Attention context (orange) and the Thinking context (blue) in thick lines. Colored violin areas and dots represent the dispersion of the RT means of each cue-validity within each context. Colored thick error bars indicate ±1 SE of the group mean, for a between-groups comparison. Black thin error bars indicate ±1 SE of the group mean, corrected to within-subject variability (Cousineau & O’Brien, 2014), for within-group comparisons.

**Figure 3.**
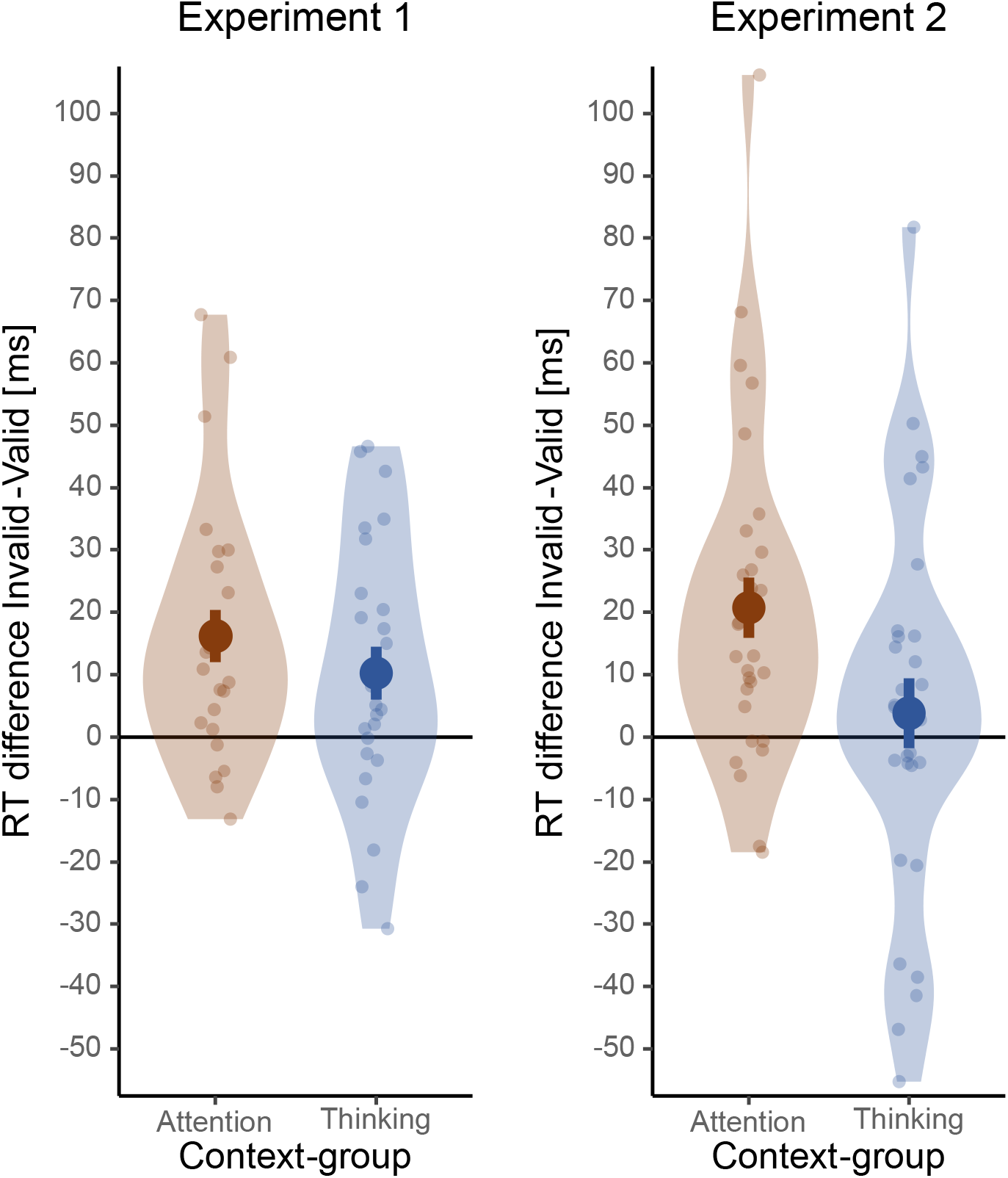
*GCE in each context-group*, in **Experiment 1** (left; N=50) and **Experiment 2** (right; N=60), presented as the mean RT difference. Individual RT differences were computed by subtracting the mean RT in valid trials from the mean RT in invalid trials, for each participant. Colored violin areas and dots represent the dispersion of individual GCEs within each context. Error bars indicate ±1 SE of the GCEs’ group mean.

These findings converge to support the research hypothesis: the GCE, which was clearly evident in the control (Attention) group, was abolished in the Thinking group. When contextual information (i.e., the task-irrelevant arithmetic problem) suggested that the observed individual was engaged in thinking rather than attending, the GCE disappeared. These results demonstrate that the GCE is neither automatic nor reflexive; rather, it is highly dependent on specific context and the top-down interpretation of the environment.

### Experiment 2

RTs were modeled similarly to **Experiment 1**, except that the model also allowed for adding the interaction between Trial number and Group as an additional covariate. As in Experiment 1, we found a significant main effect for cue-validity (χ^2^(4) = 38.98, p < 0.001), reflected by faster RTs towards validly cued targets, compared to neutral (β = 12.0ms, t = 3.14, p = 0.001), but not compared to invalidly cued targets (β = 4.4ms, t = 1.14, p = 0.25). We also found a significant main effect for context (χ^2^(4) = 17.8, p = 0.001), resulting from shorter RTs in the Attention group relative to the Thinking group (β = -88.6 ms, t = -2.52, p = 0.012). Confirming our hypothesis, we found a significant interaction between group and cue-validity (χ^2^(2) = 9.40, p = 0.009), reflected by a GCE (the difference in RT between valid and invalid trials) in the Attention group (β = 13.8ms, t = 2.69, p = 0.007). As described above, this interaction was not found to be significant in the Experiment 1, likely due to its lower statistical power. Two one-way ANOVAs conducted separately within each group were consistent with our hypothesis as they revealed a GCE only in the Attention group (F(2,7874)=10.46, p < 0.001) but not in the Thinking group (F(2,7807)=1.24, p= 0.29).

These findings were supported by a Bayesian analysis suggesting very strong evidence for a GCE in the Attention group (BF_10_=49.93) and extreme evidence for no GCE in the Thinking group (BF_10_=0.005). More specifically, In the Attention group, RTs in valid trials were found to be significantly faster compared to invalid (M_Δ_ = 20.4ms, SE = 4.63, t(7874) = 4.41, p < 0.001, Hedges’s g = 0.12; 95% CI for the effect size = [0.07, 0.18]) and compared to neutral trials (M_Δ_ = 15.0ms, SE = 4.63, t(7874) = 3.25, p = 0.002, Hedges’s g = 0.09; 95% CI for the effect size = [0.04, 0.14]). However, in the Thinking group, no significant difference was found between valid and invalid trials (p = 0.47, Hedges’s g = 0.02; 95% CI for the effect size = [-0.08, 0.03]) nor between valid and neutral trials (p = 0.35, Hedges’s g = 0.04; 95% CI for the effect size = [-0.01, 0.10]).

To summarize, Experiment 2 was consistent with the Experiment 1 as it supported the hypothesis that context and cognitive interpretation modulates the attentional effect of gaze.

## Discussion

The findings of two experiments revealed that the interpretation of another person’s eye movements is intricately tied to the context in which they are perceived. The well-established Gaze Cueing effect (GCE) was abolished when participants were led to believe that the individual conducting the gaze shift was engaged in cognitive processing, rather than attentional orienting. This indicates that when individuals are perceived as focused on internal cognitive activities, their eye movements lose significance as spatial cues for attention.

### The effect of kinematics on the interpretation of eye movements

These findings highlight the critical role of context in interpreting non-verbal social cues. Although the GCE is a robust attentional phenomenon, which can be triggered by a wide array of gaze images, including schematic, photographed or real-person gaze, we found that it can be reduced by a subtle manipulation of context. By modifying the participants’ interpretation of other person’s eye movement, the effect of eye movements as attention cues was reduced. These findings challenge the hypothesis that specific kinematics of gaze aversions (GAs), particularly their predominantly upward direction, inherently serve as a social cue for cognitive processing (Servais et al., 2022). In our study, both experimental groups observed identical eye movements, yet differing contextual interpretation led to distinct attentional effects.

While our findings illustrate the impact of context on interpreting eye movement, it is important to note the potential role of kinematics in supporting specific interpretations. The eye movements that were observed in our study were always directed diagonally and upward (left-up or right-up), to align with reported characteristics of GA in existing literature (McCarthy et al., 2008). Different directions, such as downward shifts, may lead to different interpretations: such as a weaker association with effortful thinking and a stronger association with the shifting of spatial attention. Further investigation with GAs in various directions could provide insights into the influence of kinematics on interpretation.

### Top-down effects on gaze cueing

It is widely thought that the GCE is an involuntary and a reflexive attentional effect. Evidence supporting this view include studies who compared the attentional effect of non-informative social cues to that of non-informative abstract cues (e.g. arrows). Findings show that the reflexive orienting caused by a non-informative social cue is stronger than that caused by a symbolic cue (Driver et al., 1999; Ristic, Wright & Kingstone, 2007; Schmitz et al., 2024). However, even reflexive processes can be modulated by top-down control.

Our findings revealed how high-level processes, such as interpreting the cognitive states of others, can modulate reflexive attentional mechanisms like the GCE. Other studies have previously explored top-down effects on the GCE, particularly the effect of the cueing face’s perceived ability to see the target. Findings in this area have been mixed: some studies suggest a stronger GCE when participants believe the cueing face can see the target (Morgan et al., 2018; Teufel et al., 2010), whereas others report no such effect (Cole et al., 2015; Kingstone et al., 2019). One possible explanation for these discrepancies is the variation in methodologies used to manipulate cue interpretation. For instance, Teufel et al. (2010) and Morgan et al. (2018) influenced participants’ interpretations via explicit instructions, informing them that certain glasses worn by the cueing face were opaque while others were transparent. In contrast, Cole et al. (2015) manipulated mental state attribution by altering stimulus properties, such as adding or removing occluding barriers between the gaze cue and the target. These methodological differences suggest that explicit, top-down contextual manipulations may override the reflexive cueing effect, whereas purely visual manipulations may not. Our findings align with this perspective, as the attentional modulation in our study was driven by a top-down contextual manipulation of context, rather than by changes in the visual properties of the gaze cue.

A study by Schmitz and Einhäuser (2023) aligns with our findings, showing that the interpretation of a symbolic cue can influence its attentional cueing effect, and that it can even be perceived as a social cue when contextually primed. This supports our claim that context impacts gaze cueing.

However, while this finding highlights the influence of context and interpretation on spatial cueing, it is distinct from the present findings. Specifically, their finding demonstrates that a symbolic cue can be interpreted as a social one, while the present findings show that the observation of a social cue, such as a person shifting their gaze sideways, does not invariably function as a social cue. We demonstrate here that what is typically considered a clear and direct social cue may, in fact, be perceived as a non-cue, depending on contextual factors. This distinction is subtle but crucial; while the findings by Schmitz and Einhäuser illustrated the effects of context and interpretation, they do not establish that social gaze cueing can be abolished based on interpretation, as the present findings do.

### Gaze aversions as a social cue for cognitive processing

Our results emphasize context as essential for interpreting non-verbal social cues, including GAs. It is important to note in this respect that the exact purpose of GAs during cognitive processing remains incompletely understood. While prior studies, including our own, primarily emphasized GAs’ role in minimizing distractions and conserving cognitive resources (Doherty-Sneddon & Phelps, 2005; Abeles & Yuval-Greenberg, 2017, 2021), several studies suggested that GAs serve also social functions (Andrist et al., 2014; Doherty-Sneddon et al., 2005; Ho et al., 2015; McCarthy et al., 2008). However, this previous research examined GAs primarily from the perspective of the individual exhibiting them, focusing on the triggers for these aversions, modulation, and suppression. The present findings offer a novel approach as they investigate GAs, for the first time, the perspective of the observers of these eye movements. Namely, they reveal that people who observe GAs in other people interpret them based on their context, in ways that can influence their function as attention cues.

We propose that a fundamental social function of GAs is to signal ongoing cognitive processing, allowing observers to recognize when an individual is engaged in complex mental tasks. This interpretation aligns with the idea that GAs can serve as implicit cues prompting interlocutors to reduce conversational input, slow down or pause until cognitive processing is complete. Supporting this hypothesis, research has found that GA amplitude differs between social and non-social settings. In non-social contexts, GAs are typically subtle with minimal amplitude of around 1° (Abeles & Yuval-Greenberg, 2021), making them less conspicuous to external observers (Gibson & Pick, 1963). In contrast, in social environments, GAs become more pronounced (unpublished data), potentially reflecting an adaptive mechanism for signaling cognitive engagement. Future research could systematically manipulate social and cognitive demands to determine how these variables influence GA characteristics, including direction and amplitude.

### Main effect of context

Beyond the anticipated contextual influence on attention, the findings also revealed an unexpected main effect of context, wherein RTs were generally longer in the Thinking group compared to the Attention group. One potential explanation for this effect is that participants in the Thinking group were inadvertently influenced by the auditory math questions presented during each trial. Despite being instructed to ignore the arithmetic content, some participants in this group may have engaged in mental calculation, increasing their cognitive load and slowing response times. An alternative explanation is the slightly longer duration of experimental sessions in the Thinking group due to the additional answering video in each trial. This could have led to increased fatigue, contributing to longer reactions. However, this fatigue-related explanation was ruled-out, as there was no evidence of progressive slowing over the course of the session (see Supplementary Material S1). Regardless of the cause, this context effect does not confound the study’s key findings, as it affected all three attentional conditions equally.

### Constraints on Generality

The present study provides initial evidence that the interpretation of gaze critically influences the gaze cueing effect (GCE). However, several methodological decisions limit the generalizability of these findings. We examined this effect using a single male actor, restricting cognitive processing to arithmetic calculation, and employing a basic perceptual letter discrimination task. These constraints leave open the question of whether our findings generalize beyond this specific context. Future research should test a broader range of actors, cognitive tasks, and perceptual demands to robustly establish the scope and external validity of this context-dependent modulation of the GCE.

## Conclusion

This study shows that the well-known Gaze Cueing effect (GCE) is not purely reflexive but is instead subject to top-down modulation based on contextual interpretation. When individuals believe that gaze shifts are linked to cognitive processing rather than attentional orienting, the typical GCE is attenuated. This finding underscores the fundamental role of context in shaping perception and utilization of non-verbal social cues. Furthermore, the findings demonstrate that social-cognitive mechanisms can override automatic attentional responses, revealing the dynamic interplay between high-level cognition and low-level perceptual processes. By illustrating how subtle contextual shifts can reshape gaze-based attentional effects, this study offers a new insight into the flexibility of social attention and its integration with cognitive interpretation.

