## Supplementary Material S1 for "When a look means nothing: contextual interpretation abolishes the Gaze Cueing Effect"

### Supplementary Materials

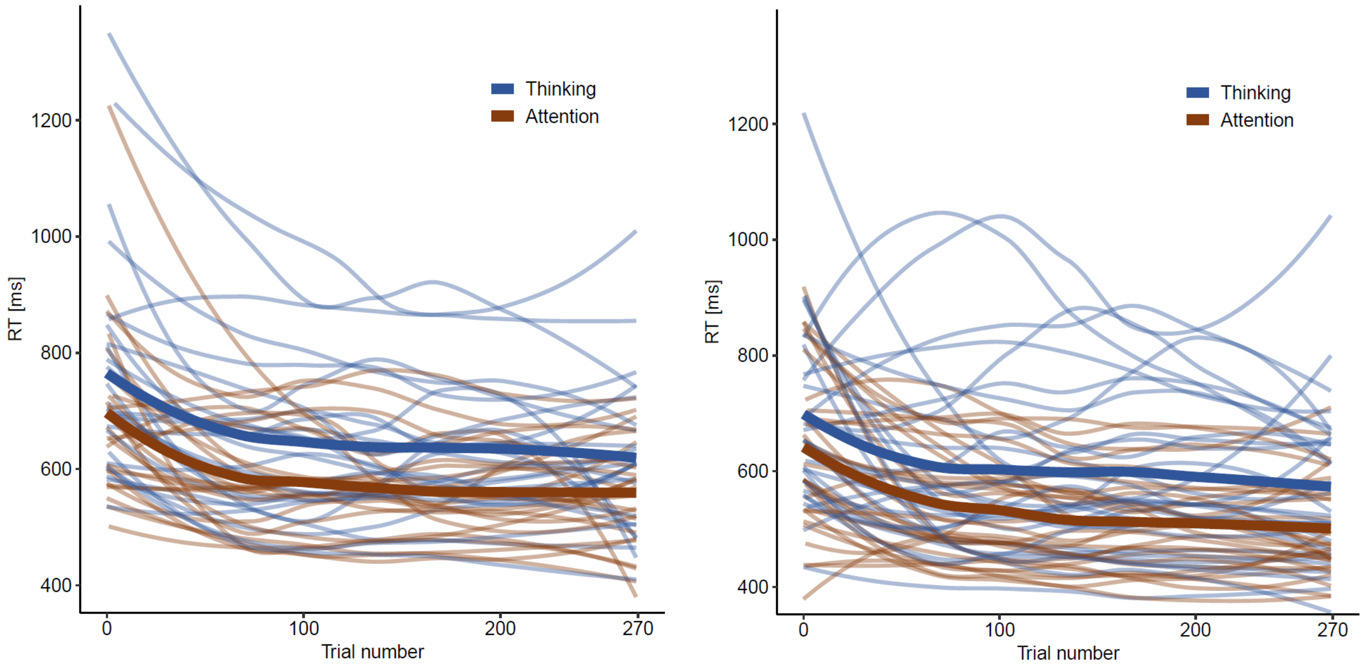


*Figure S1*. RT as a function of trial number in **Experiment 1** (left) and **Experiment 2** (right), for individuals in the Thinking group (blue) and the Attention group (brown). A smoothed local-regression interpolation of the RT carried out using the locally estimated scatterplot smoothing (LOESS) function in the R stats package (Cleveland & Loader, 1996). Individuals and group-averaged curves represented by faded-thin and thick lines, respectively.
